# Circadian Modulation of Excitation—Inhibition Balance Drives Ictal Transitions in a Mechanistic Model of Epileptic Networks

**DOI:** 10.64898/2026.05.22.726242

**Authors:** Nesrine Dlima, Ilaria Carannante, Alain Destexhe, Viktor Jirsa, Mohamed Hedi Bedoui, Damien Depannemaecker

## Abstract

Epileptic seizures emerge from pathological synchronization in neuronal networks and are strongly influenced by circadian rhythms. Here, we developed a computational framework to investigate how circadian modulation of excitation/inhibition (E/I) balance shapes transitions from physiological to pathological activity. The model consists of interacting excitatory and inhibitory populations containing varying proportions of impaired neurons with altered intrinsic excitability. Circadian effects were incorporated through modulation of synaptic time constants, mimicking daily fluctuations in E/I dynamics. Network activity was characterized using firing rates and the Spike Time Tiling Coefficient (STTC), enabling simultaneous assessment of excitability and synchrony. Our results show that seizure-like dynamics arise from nonlinear interactions between network composition, neuronal impairment, and synaptic kinetics. Distinct dynamic regimes emerged, separated by sharp transitions in synchrony and activity patterns. These findings provide a mechanistic link between circadian regulation and seizure susceptibility, supporting the development of chronotherapy approaches for epilepsy.

## 1 Introduction

Epilepsy affects more than 50 million people worldwide and is characterized by recurrent, unprovoked seizures arising from pathological hypersynchronization in neuronal networks. These events are not random; instead, seizure occurrence follows robust temporal patterns shaped by endogenous physiological rhythms, particularly sleep–wake cycles and circadian oscillations. This temporal structure highlights circadian modulation of brain excitability as a key component of epilepsy pathophysiology [1–4].

At the neurobiological level, circadian rhythms regulate neuronal function across multiple scales, from gene expression to large-scale network dynamics. A substantial proportion of genes involved in synaptic transmission exhibit circadian oscillations, directly influencing excitatory and inhibitory synaptic properties [5, 6]. These molecular fluctuations translate into changes in glutamatergic and GABAergic receptor kinetics, as well as alterations in functional connectivity within cortical networks. In a previous study, the importance of circadian modulation to reshape neuronal excitability and alter seizure susceptibility at the level of individual neurons has been shown [7–9]. These findings provided mechanistic evidence suggesting that slow physiological rhythms continuously modulate the intrinsic stability of neuronal states. However, epileptic seizures do not arise solely from isolated neuronal dysfunction. Rather, they emerge from collective interactions involving large-scale synchronization, pathological recruitment, and dynamic reorganization of neuronal populations [10, 11]. Experimental studies in both animals and humans, including transcranial magnetic stimulation combined with EEG, demonstrate that the excitation/inhibition (E/I) balance is dynamically modulated across the circadian cycle [12, 13].

In epilepsy, disruption of the E/I balance is a central mechanism underlying seizure generation. Electrophysiological recordings have identified distinct seizure onset patterns, including hypersynchronous onsets associated with reduced inhibition or increased excitation, and low-voltage fast activity involving more complex interactions between excitatory and inhibitory populations [14]. Despite these advances, the mechanisms by which circadian fluctuations in E/I balance interact with network dynamics to trigger transitions from interictal to ictal states remain poorly understood. In particular, bridging molecular and cellular processes with emergent network properties such as synchrony and seizure propagation remains a major challenge.

To address this gap, we developed a computational model building to investigate how pathological activity emerges from nonlinear interactions within neuronal networks under varying levels of impairment. We focus on alterations in E/I balance, a primary target of most antiepileptic drugs, and propose a mechanistic framework to ultimately support chronotherapy strategies, where treatment timing is aligned with individual circadian rhythms.

Specifically, we examine seizure emergence in networks containing impaired neurons embedded within interacting excitatory and inhibitory populations. Impairment is modeled based on previous work [15] and systematically varied in both severity and proportion within the network. We further incorporate circadian modulation through temporal variations in synaptic time constants, allowing us to study how daily rhythmic changes influence network stability and seizure susceptibility.

Characterizing the resulting network dynamics requires capturing both global activity levels and fine-scale temporal coordination. While firing rates provide a measure of overall network excitability, they do not fully describe the temporal organization of neuronal interactions. Therefore, we complement rate-based metrics with measures of synchrony. Neural synchrony is a hallmark of ictal states, reflecting coordinated activity beyond simple increases in firing rate. The Spike Time Tiling Coefficient (STTC) provides a robust and bias-resistant measure of spike train correlation, independent of firing rate [16, 17]. Combining these measures enables a more comprehensive characterization of transitions between interictal and ictal dynamics.

Our results demonstrate that the transition from physiological to pathological activity emerges from the interplay between network composition, neuronal excitability, and modulation of synaptic kinetics as observed in circadian fluctuations.

## 2 Results

The transition from physiological to pathological activity emerges from the complex interplay between network composition, neuronal excitability, and synaptic kinetics.

To investigate these mechanisms, we simulated large-scale spiking neural networks (10 000 neurons) composed of inhibitory fast-spiking neurons (FS, 2000) and excitatory regular-spiking (RS) and impaired (IMP) neuronal populations (8000, see Methods and Figure 1 for details). IMP neurons represent pathologically altered excitatory neurons with epileptic-like behaviour. FS and RS neurons were modelled using the Adaptive Exponential Integrate-and-Fire (AdEx) formalism, while IMP neurons were modelled using an extended AdEx formulation (Figure 2). The latter includes a slow modulatory variable associated with ionic dysregulation and altered intrinsic excitability. Within this framework, the parameter *Z*_0_ controls the excitability regime of IMP neurons and phenomenologically captures shifts in extracellular potassium dynamics (see Methods for details).

**Fig. 1.**
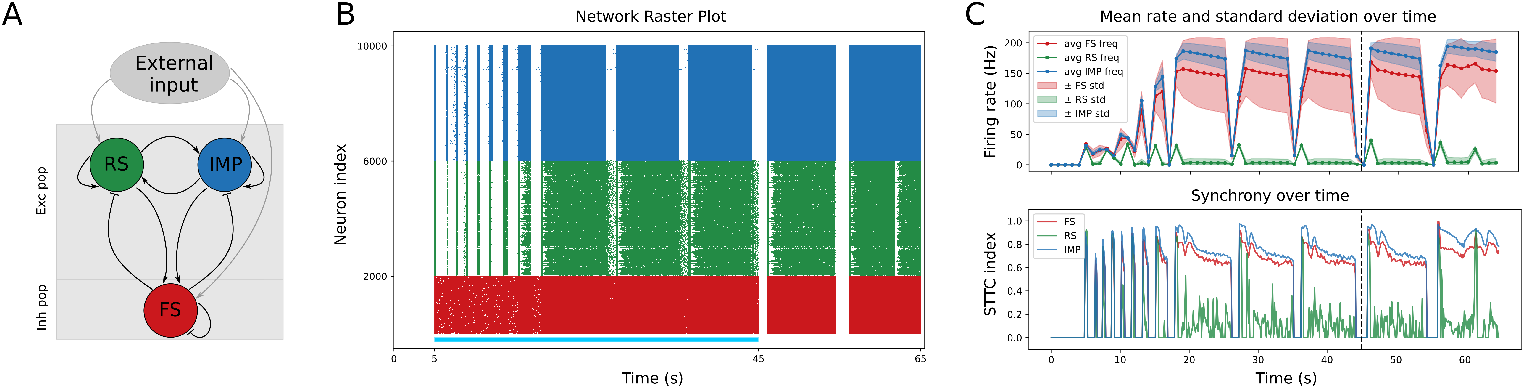
Network composition and measures. A) Schematic of the network architecture, comprising an inhibitory population of Fast Spiking (FS, red) and two excitatory populations: Regular Spiking (RS, green) and Impaired neurons (IMP, blue). The populations are all-to-all connected with 5% probability. Each receives external stimulation driving the system. Networks consist of 10 000 neurons, 20% FS and the remaining 80% is divided between RS and IMP neurons in varying proportions. B) Representative network raster plot where IMP neurons comprise 50% of the excitatory population (4000 RS and 4000 IMP). Simulations include a 5-second transient period, external input from 5 to 45 seconds (indicated by the light blue horizontal bar), and a 20-second recovery phase. C) Quantitative analysis of network dynamics, showing the mean firing rate and standard deviation over time (top) and the Spike Time Tiling Coefficient (STTC) as a measure of synchrony over time (bottom). Unless otherwise specified, the *Z*_0_ value for the IMP models is set to *−*50mV, and the ratio between inhibitory and excitatory synaptic time constants (Tau ratio) is 1.

**Fig. 2.**
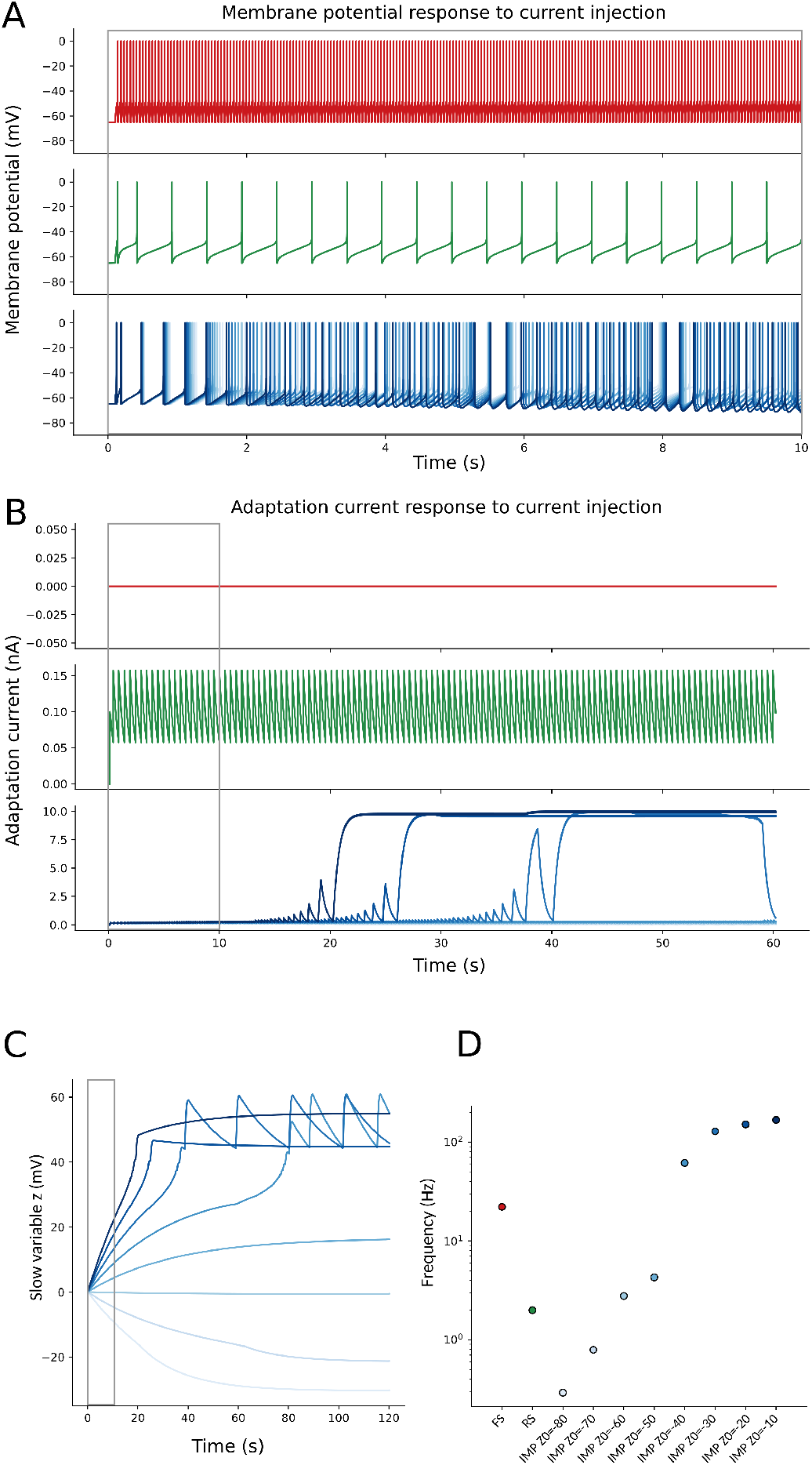
Single-neuron responses to current injection across neuron types and impairment levels. A) Membrane potential responses. FS neuron (red) displays sustained high-frequency firing, RS neuron (green) exhibits regular spiking, and IMP neurons (blue shades) show progressively increased firing rates and altered spike patterns as *Z*_0_ increases (from *−* 80mV to *−* 10mV), reflecting enhanced excitability. B) Adaptation current dynamics. While FS exhibits no adaptation, RS shows regular adaptation oscillations and IMP neurons display slow, cumulative changes in the adaptation current. C) Time evolution of the slow variable *z* in IMP neurons. Increasing *Z*_0_ leads to higher steady-state values of *z*, providing a slow excitability drive that promotes sustained and high-frequency firing regimes. D) Output firing frequency across neuron types and impairment levels. IMP neurons show monotonic increase in firing rate with increasing *Z*_0_. Grey boxes across panels A-C highlight the initial 10 seconds of stimulations. Simulations are performed on single, uncoupled neurons under identical current injection (0.2nA), while the displayed time windows differ across panels to best capture the different dynamics.

To characterize the landscape of the network dynamics, we performed a systematic exploration across three primary control parameters: the proportion of impaired excitatory neurons (*IMP percentage*), the level of intrinsic excitability (*Z*_0_), and the ratio of inhibitory-to-excitatory synaptic decay times (*Tau ratio*). This allowed us to evaluate how circadian modulation of the inhibition/excitation balance might drive transitions between network states.

By integrating population firing rates with rate-independent synchrony measure (STTC, see Methods for details) and temporal classification of network states, we demonstrate that the system undergoes regime shifts.

### 2.1 Increasing pathological recruitment drives the network towards pathological states

To investigate how pathological recruitment reshapes network activity, we systematically varied the proportion of IMP neurons while keeping their intrinsic excitability fixed (*Z*_0_ = *−*50mV). In all simulations, the inhibitory FS population remained constant (2000 neurons), while the excitatory population was progressively shifted from RS to IMP. For example, 10% IMP pop corresponds to 800 IMP and 7200 RS, while 50% and 100% IMP pop correspond to equal RS/IMP populations and complete pathological recruitment of excitatory population, respectively.

As the proportion of IMP neurons increased, the network underwent a progressive reorganization of its dynamical state (Figure 3). At low IMP fractions (10%), activity remained sparse, weakly synchronized, and predominantly interictal (Figure 3, top row). Increasing the pathological recruitment to intermediate levels (50%) produced intermittent transitions toward highly active and synchronized episodes separated by periods of lower activity (Figure 3, central row). At full impairment (100%), the network rapidly reached a persistent high-activity regime characterized by sustained synchrony and prolonged ictal occupancy. Importantly, the transition was not limited to a simple increase in the firing rate (Figure 3A). Higher IMP fractions also altered the temporal structure of network activity, leading to longer-lasting synchronized episodes (Figure 3B). Together, these results indicate that increasing the proportion of IMP neurons progressively destabilizes physiological activity and promotes the emergence of persistent pathological network states.

**Fig. 3.**
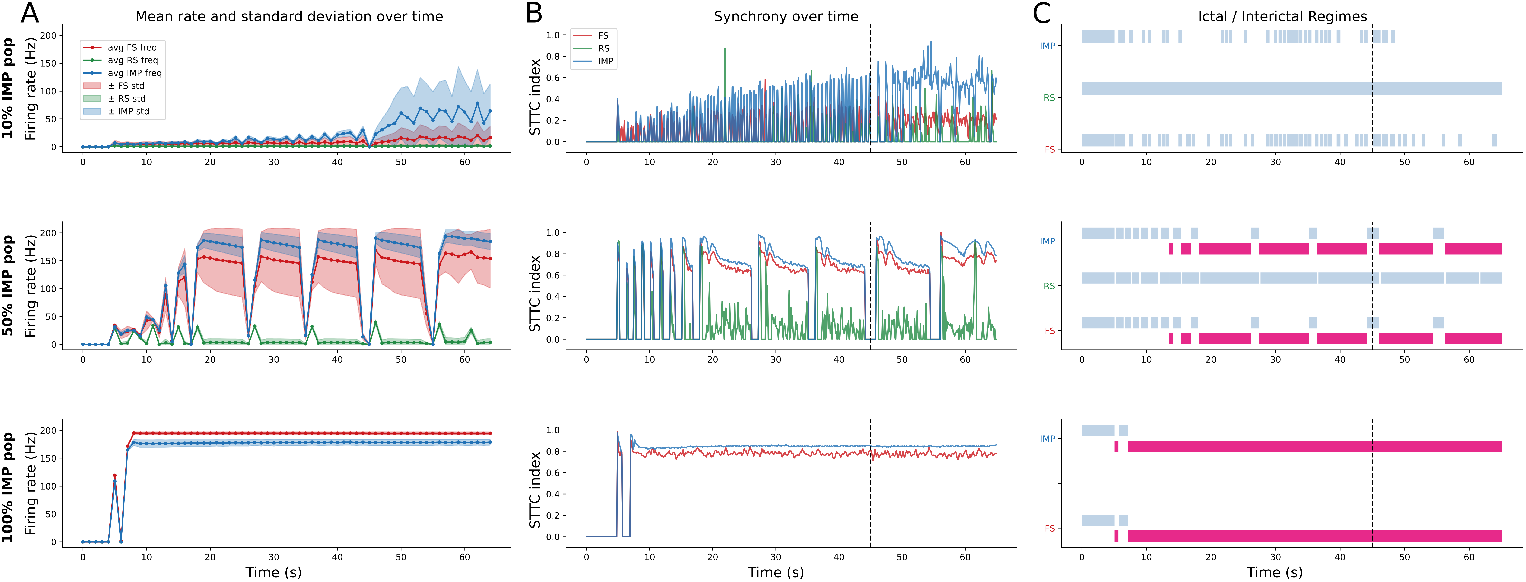
Impact of impaired neuron proportion on network dynamics and ictal transitions. Mean firing rate (solid lines) and standard deviation (shaded areas) over time for increasing proportion of impaired (IMP) neurons within the excitatory population (top to bottom: 10%, 50%, 100%). At low IMP fractions, activity remains sparse and irregular. At intermediate levels, the network exhibits recurrent high-frequency activities separated by short quiescent periods. At full impairment, the system rapidly transitions to a sustained high-rate regime. B) Synchrony dynamics is quantified using the Spike Time Tiling Coefficient (STTC). Low IMP fraction produces weak and fluctuating synchrony. At high IMP fractions, synchrony saturates (central panel) and remains persistently elevated (bottom panel), indicating a stable pathological state. The dashed vertical line marks the end of the external stimulation. C) Temporal classification of network states into ictal (magenta) and interictal (light blue) regimes based on firing rate and synchrony thresholds. Increasing IMP proportion leads to longer and more frequent ictal periods.

### 2.2 Intrinsic excitability and pathological recruitment jointly shape ictal regimes

After establishing that increased pathological recruitment destabilizes network activity (Figure 3), we next investigated how intrinsic excitability and network composition jointly shape ictal dynamics of the IMP population. To this end, we systematically explored the two-dimensional parameter space defined by the intrinsic excitability parameter (*Z*_0_) and the proportion of IMP neurons, and quantified the resulting activity measuring the number of ictal events (Figure 4A), their duration (Figure 4B) and the synchrony of the IMP population (Figure 4C). The network dynamics revealed a strong interaction between excitability and pathological recruitment, leading to the emergence of distinct dynamical regimes rather than simple monotonic progression toward pathological activity. As either the intrinsic excitability or pathological recruitment increased, the temporal organization of the activity changed substantially. Intermediate parameter regimes were characterized by frequent transient ictal episodes, while stronger excitability (greater values of *Z*_0_) produced fewer but longer events (Figure 4A, B). Indeed, the reduction in seizure like events at high *Z*_0_ did not reflect a recovery toward physiological dynamics, but progressively prolonged pathological states occupying a larger fraction of the simulation. This transition toward persistent activity was accompanied by a marked increase in network synchrony (Figure 4C). STTC values increased with *Z*_0_ across all IMP fractions, but the effect became substantially stronger as the proportion of IMP neurons increased. Together, these results reveal a trade-off between ictal event frequency and persistence within the IMP population.

**Fig. 4.**
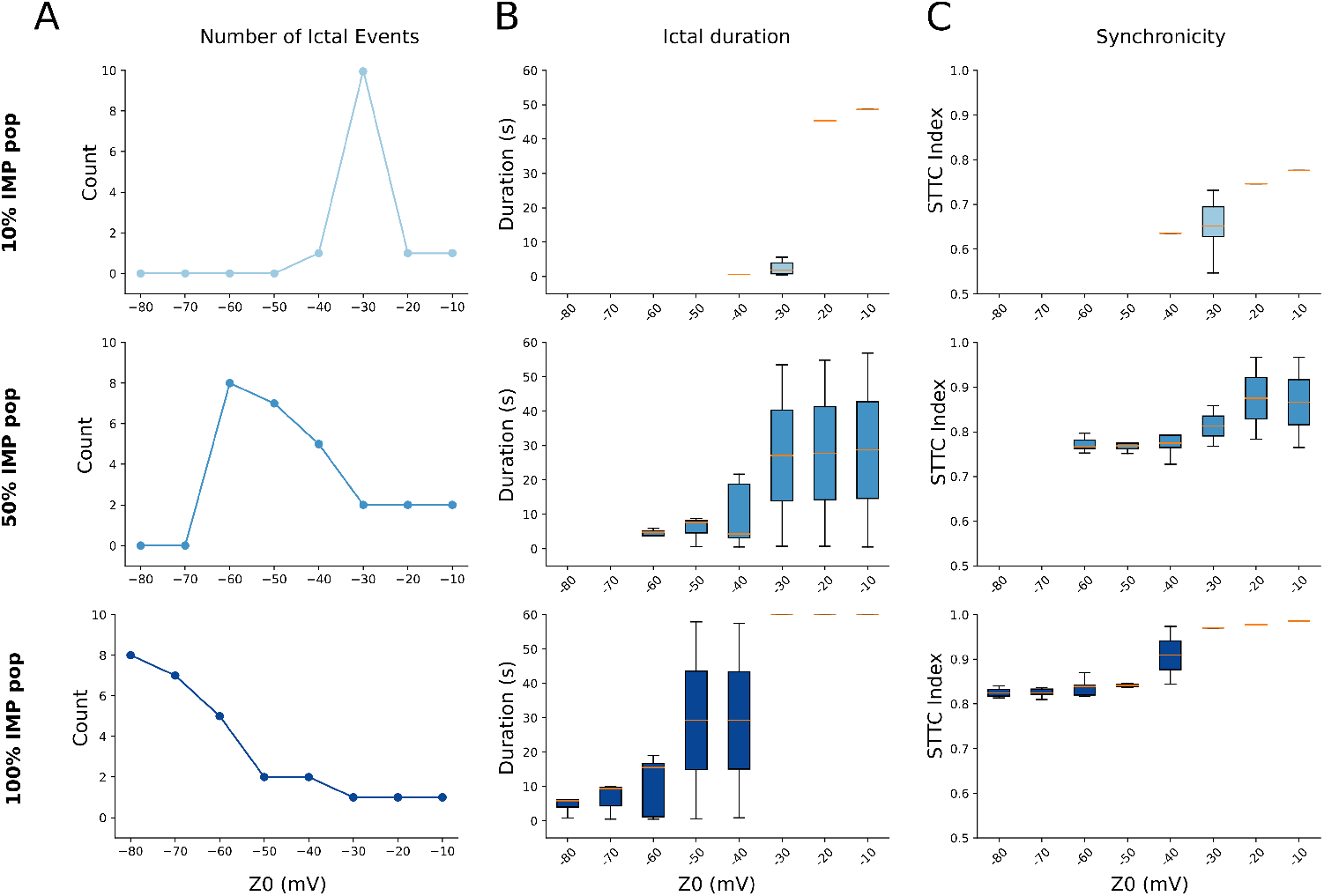
Effect of *Z*_0_ on ictal dynamics across network compositions. A) Number of ictal events in the IMP population as a function of *Z*_0_, for increasing proportions of IMP neurons (top to bottom: 10%, 50%, 100%). At low IMP fraction, ictal events emerge only within a narrow range of *Z*_0_ values. At higher IMP fractions, events are present across a broader range but progressively decrease in number as *Z*_0_ increases. This is because their duration increases, as shown in B), where the distribution of ictal event duration is represented. Increasing *Z*_0_ leads to longer-lasting events, with a marked shift from brief, to prolonged episodes. At high IMP fraction, the durations saturate, consistent with the emergence of sustained pathological activity. C) Network synchrony quantified by the Spike Time Tiling Coefficient (STTC) and as a function of *Z*_0_. Synchrony increases with *Z*_0_ across all conditions, reaching its highest levels at 100% IMP fraction. These results reveal a compromise between ictal event frequency and duration, with increasing excitability (via increase in *Z*_0_) promoting persistent pathological activity. .

### 2.3 Synaptic timescale modulation influences ictal persistence and network synchrony

We next investigated how modulation of the inhibitory-to-excitatory synaptic decay time ratio (Tau ratio) reshapes network synchrony and ictal persistence in both FS and IMP populations (Figure 5). Because circadian rhythm is known to influence excitation/inhibition balance, the Tau ratio was used here as a phenomenological parameter to mimic circadian synaptic modulation.

**Fig. 5.**
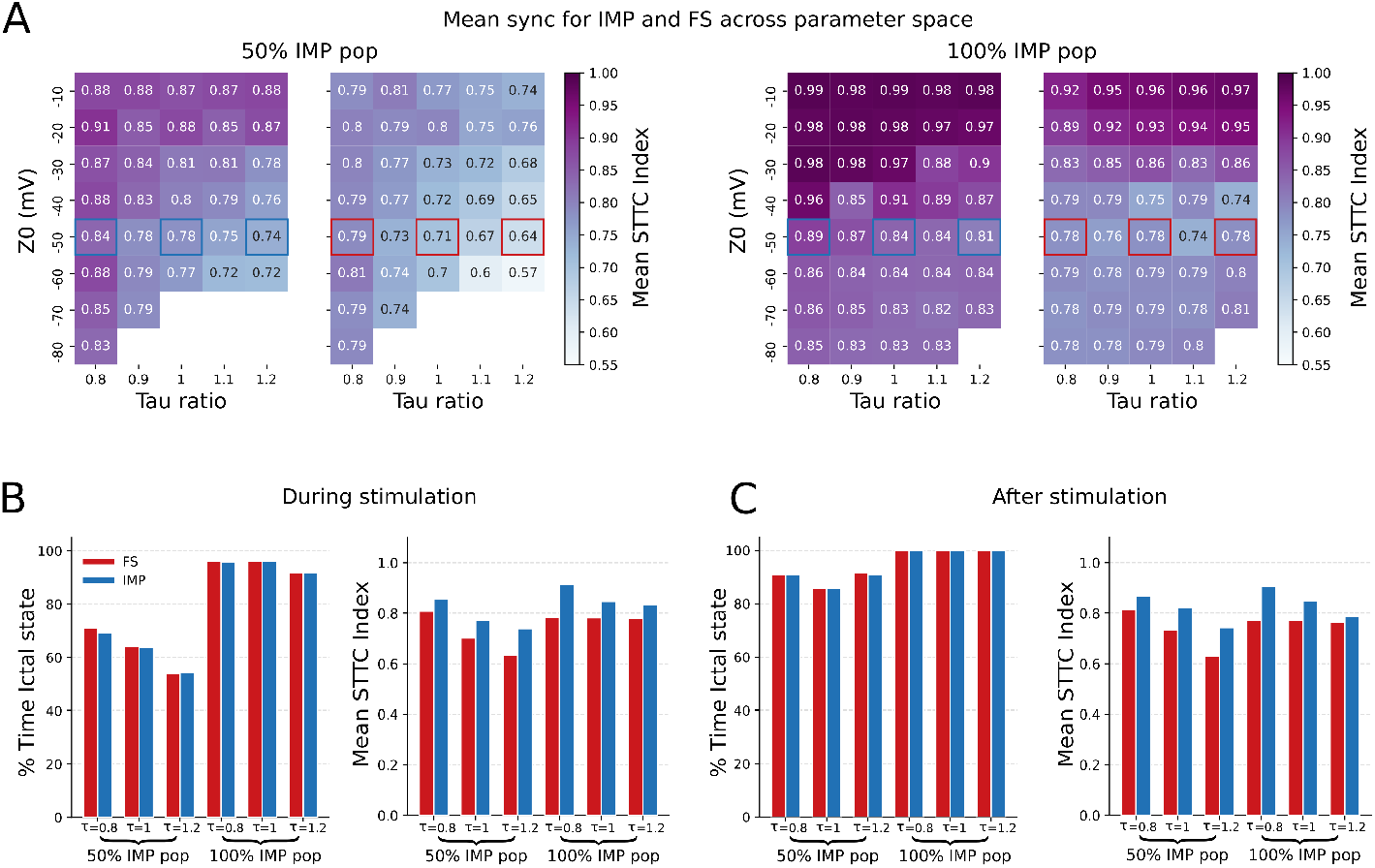
Effect of synaptic time-scale ratio on ictal synchrony in IMP and FS populations. A) Mean Spike Time Tiling Coefficient (STTC), computed during ictal phases, for IMP (left subpanels) and FS (right subpanels) population across parameter space defined by *Z*_0_ (y-axis) and the synaptic time-scale ratio (Tau ratio as in eq. 6). The Tau ratio controls the relative duration of the decay of inhibitory and excitatory synaptic time constants. The results are shown for two levels of epileptic recruitment: 50% (left) and 100% (right) of IMP neurons, corresponding to 4000 and 8000 IMP neurons, respectively (and hence 4000 and 0 RS neurons, respectively). Color indicates the mean STTC, and values are reported in each square. Highlighted squares indicate representative parameter combinations used for the more detailed analyses in panels B and C. B) During stimulation (5-45 s), percentage of time spent in the ictal state (left) and corresponding mean STTC during ictal periods (right) for FS (red) and IMP (blue) populations. Results are shown for selected values of the Tau ratio (0.8, 1 and 1.2) and for two levels of IMP recruitment (50% and 100%). C) Same as B) but computed after stimulation (45-65 s), capturing the persistence and reorganization of ictal dynamics following the transient input.

Both FS and IMP populations exhibited similar large-scale dynamical organization despite quantitative differences in synchrony levels (see also Supplementary Figure 2 and 3). IMP populations generally reached higher STTC values, particularly in strongly pathological regimes. However, transitions between low to high synchrony regimes occurred in comparable regions of parameter space. Nevertheless, this correspondence depended strongly on the proportion of impaired neurons. At low IMP fractions (10 *−* 20%), the IMP population could transiently enter ictal-like synchronized states while the FS population largely remained outside the ictal regime. As the proportion of IMP neurons increased, both populations progressively converged toward similar synchrony landscapers (Supplementary Figure 2 and 3).

The Tau ratio shaped these transitions. Lower Tau ratio values, corresponding to faster inhibitory decay relative to excitation, promoted highly synchronized states and expanded the parameter regions associated with pathological activity (Figure 5A). Intermediate pathological recruitment (50% IMP) revealed regions of high sensitivity, where relatively small changes in synaptic timescales substantially altered synchrony (Figure 5A left panels, gradient of decreasing STTC indexes moving along the rows). In contrast, at high IMP fractions, the network converged toward stable hypersynchronous regimes over a broad range of parameters and the Tau ratio had less impact (Figure 5A right panels).

To further characterize this effect, we separately quantified network dynamics during stimulation and after stimulation offset (see Methods for details). During stimulation, both populations showed prolonged ictal occupancy and increased synchrony as the Tau ratio decreased. Importantly, the post-stimulation dynamics clearly revealed that the networks failed to recover toward low-activity states after the external drive terminated. Instead persistent ictal activity and elevated synchrony remained sustained throughout the recovery period (period without external stimulation). The 50% IMP condition retained a strong dependence on synaptic timescales even after stimulation offset, suggesting that intermediate pathological recruitment preserves degree of dynamical flexibility and recoverability.

Together these results identify synaptic timescale modulation, and hence circadian-like synaptic modulation, as a control mechanism governing not only the emergence of ictal activity, but also its persistence and recovery dynamics.

### 2.4 Organization of dynamic regimes in parameter space

To obtain a global view of how pathological recruitment (IMP fraction), intrinsic excitability (*Z*_0_) and synaptic timescale (Tau ratio) organize network behaviour, we systematically mapped the IMP population synchrony landscape across *Z*_0_ and Tau ratio for increasing fractions of IMP neurons (Figure 6 and Supplementary Figure 2).

**Fig. 6.**
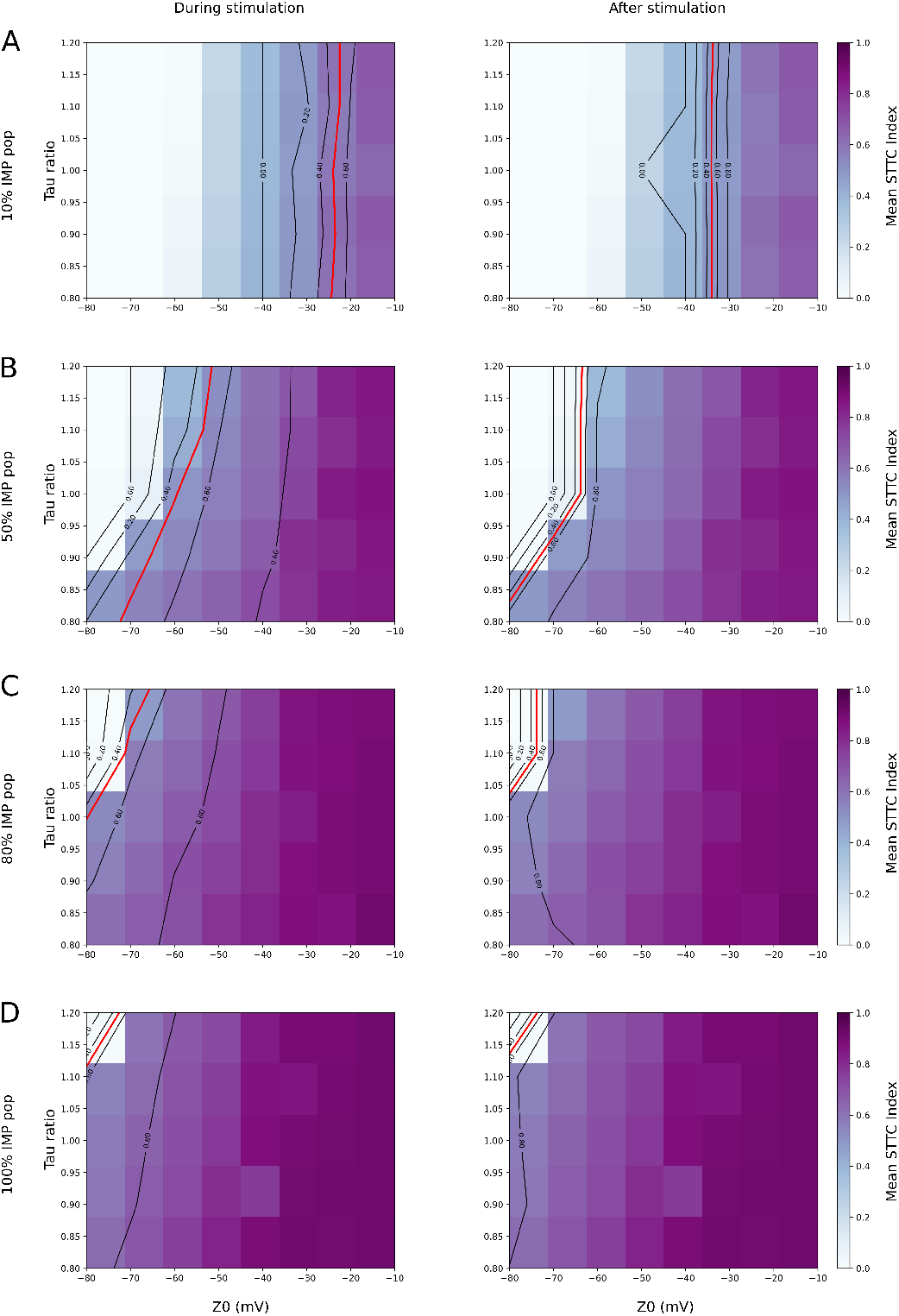
Synchrony landscape across parameter space and impaired population fraction in IMP population. Heatmaps show the mean Spike Time Tiling Coefficient (STTC), computed over IMP populations, as a function of the parameter *Z*_0_ and the synaptic time-scale ratio (Tau ratio as in eq. 6), respectively on the x and y axis. Rows correspond to increasing fractions of impaired neurons (A: 10%, B: 50%, C: 80%, D: 100%) Columns represent different temporal windows: during stimulation (left, 5 - 45 s) and after stimulation (right, 45 - 65 s). Color indicates level of synchrony (mean STTC), black contour lines denote iso-levels of synchrony, with the red line highlighting the transition boundary (threshold=0.5) separating low- and high-synchrony regimes. Across all conditions, increasing *Z*_0_ leads to higher synchrony, but the structure of the transition depends strongly on the fraction of impaired neurons. At low IMP fractions (A), synchrony remains confined to a narrow region of parameter space with Tau ratio having just a minimal influence. At higher fractions (B-D) the high-synchrony regime expands, becoming more structured and is strongly modulated by the Tau ratio. Lower values (faster inhibitory dynamics compared to excitation) promote the emergence of synchrony at more hyperpolarized *Z*_0_, effectively shifting the transition boundary. Comparing stimulation and post-stimulation periods reveals how network dynamics evolve once the transient excitatory input stops, highlighting the persistence of synchronous activity in the absence of sustained drive.

Across all conditions, the parameter space is organized into well-defined dynamical domains characterized by distinct levels of synchrony. These domains were separated by sharp transition regions highlighted by STTC contour lines. The resulting maps reveal that pathological recruitment progressively reshapes the topology of the dynamical landscape, expanding the regions associated with persistent synchronized activity.

At low IMP fraction (Figure 6A), synchrony remained relatively weak across most of the parameter space, and transitions between regions were gradual. Moreover, in this regime the network dynamics were only moderately sensitive to changes in intrinsic excitability or synaptic timescales, and highly synchronized states remained confined to restricted parameter regions.

Increasing the fraction of IMP neurons substantially altered this organization (Figure 6B-D). Indeed, high synchrony regimes progressively expanded toward lower values of, indicating that pathological recruitment reduced the excitability threshold required for the emergence of synchronized pathological activity. At intermediate to high IMP fraction (Figure 6B, C), relatively small variation in either *Z*_0_ or the Tau ratio produced abrupt transitions between weakly to synchronized and highly synchronized states. These transitions were reflected by increased curvature of STTC isolines, revealing the presence of sharp dynamical boundaries and enhanced susceptibility to modulation of the E-I balance.

The Tau ratio exerted a particularly strong influence in networks with a medium to high fraction of IMP population. Lower Tau ratio values, corresponding to faster inhibitory decay relative to excitation, systematically promoted the expansion of synchronized regimes and shifted the critical transition boundary toward more hyperpolatized *Z*_0_ values. Thus, circadian-like modulation of synaptic timescales effectively altered the accessibility of pathological states within the parameter space.

Importantly, the overall organization of the dynamical landscape persisted after stimulation offset (Figure 6, right panels), and the transition boundaries became closer to each other compared to the “during stimulation” counterparts (Figure 6, left panels).

Corresponding analyses for additional IMP fractions and FS populations are provided in the Supplementary Material. Together, these results reveal a progressive restructuring of the network landscape. Pathological recruitment expands the domain of synchronized activity, shifts the transition boundaries toward lower *Z*_0_, and enhances the sensitivity to synaptic modulation.

Overall, this organization of the parameter space suggests that slow modulation of the E-I balance (modelled via the Tau ratio), can progressively shift the network across critical dynamical boundaries separating physiological and pathological regimes.

## 3 Discussion

In this study, we show that ictal transitions do not arise from a simple increase in neuronal excitability, but from a multiscale reorganization of network dynamics driven by the combined effects of pathological recruitment, intrinsic excitability, and synaptic time scales. Across parameter space, the network is organized into distinct dynamical regimes separated by sharp boundaries, indicating that seizure emergence is better understood as a transition between metastable states than as a linear increase in activity. In this framework, epilepsy appears not as a fixed condition, but as a dynamic trajectory shaped by excitation-inhibition interactions and network composition.

Increasing the fraction of impaired neurons progressively destabilized the network and shifted it from irregular, weakly synchronized interictal activity toward transient and then persistent ictal regimes. At low levels of pathological recruitment, the network remained predominantly interictal, whereas intermediate and high levels of impairment promoted recurrent synchronization and prolonged seizure-like episodes. Importantly, this progression was not captured by firing rate alone. As excitability increased, the number of discrete ictal events could decrease while their duration and synchrony increased, showing that seizure burden is defined not only by event frequency but also by persistence and temporal organization.

A second major finding is that intrinsic excitability and pathological recruitment interact nonlinearly. The effect of *Z*_0_ depended strongly on the proportion of impaired neurons, with intermediate values favoring brief and intermittent ictal episodes and higher values promoting long-lasting synchronized states. This trade-off between event number and event duration suggests a qualitative change in collective dynamics rather than a simple amplification of neuronal activity. STTC analyses supported this interpretation by showing that ictal-like activity was consistently associated with stronger temporal coordination across neurons, confirming that hypersynchrony is a hallmark of the transition to persistent pathological states. These results are consistent with experimental observations indicating that seizures emerge through progressive recruitment and coordination of distributed neuronal populations [11].

The phase maps further revealed that small variations in intrinsic excitability or in the inhibitory-to-excitatory synaptic time-scale ratio could trigger abrupt transitions between low-synchrony and high-synchrony regimes. As pathological recruitment increased, the synchrony landscape became more structured and the critical boundaries became sharper, indicating greater sensitivity of the system to parametric perturbations. This organization is consistent with dynamical-systems views of epilepsy, in which seizure onset reflects proximity to critical boundaries in state space rather than a single causal trigger [18–20]. It also supports the broader view that epilepsy is a network disorder in which local pathology reshapes global stability rather than merely increasing excitability in isolation [21, 22].

These findings are particularly relevant in the context of circadian modulation. Experimental studies have shown that cortical excitability, inhibitory efficacy, and synaptic properties fluctuate across the sleep-wake cycle and circadian rhythm [2, 3, 23]. In our model, the inhibitory-to-excitatory synaptic time-scale ratio acted as a slow control parameter capable of shifting the network toward or away from critical boundaries. This suggests that circadian fluctuations may not directly trigger seizures, but instead modulate seizure susceptibility by changing the system’s position in parameter space. When the network is already close to a transition boundary, even modest physiological changes in synaptic timing may be sufficient to promote hypersynchronous activity.

The behaviour of the FS inhibitory population further illustrates how ictal dynamics emerge from collective network interactions rather than from isolated excitatory mechanisms. As the proportion of IMP excitatory neurons increased, the FS population underwent similar transitions to those in the IMP population.This behavior in the model is consistent with the possibility that strong excitatory recruitment from the IMP population, acting on FS neurons with intrinsically weaker adaptation than RS neurons, contributes to this phenomenon. As excitation increases, this reduced adaptation may predispose FS neurons to participation in hypersynchronous network states. These results are consistent with experimental observations showing that inhibitory interneurons can remain highly active during seizures and participate in the temporal organization of ictal activity [24].

A major strength of this study is the integration of network composition, intrinsic excitability, synaptic kinetics, and synchrony into a single mechanistic framework. By combining firing rate, temporal classification, and STTC, we captured both the intensity and the organization of network activity, which is essential for distinguishing brief seizure-like events from sustained pathological states. The systematic exploration of three-dimensional parameter spaces also allowed us to identify sharp dynamical boundaries and to show how these boundaries shift with pathological recruitment. These sharp transitions are consistent with observations at multiple scales in seizures [21]. This makes the model useful for interpreting how slow physiological modulation may interact with the underlying disease state to shape seizure susceptibility.

However, this work also has several limitations. First, the model remains phenomenological and does not explicitly incorporate all cellular and molecular mechanisms involved in epilepsy, such as detailed ion homeostasis, astrocytic buffering, neuromodulation, or heterogeneous microcircuit architecture. Second, circadian modulation is represented indirectly through synaptic time constants, which capture an important aspect of temporal regulation but do not fully reproduce the complexity of real circadian biology. Third, the network is highly simplified relative to clinical epilepsy, and direct quantitative mapping to patient-specific seizure dynamics will require further validation. Finally, while STTC is robust to firing-rate confounds, additional metrics of phase relationships, propagation, and spatial organization would help refine the characterization of ictal states.

Future works should extend this framework in several directions. A first step would be to couple the present network model to explicit representations of ion concentration dynamics, glial buffering, and neuromodulatory control to better approximate the biological mechanisms underlying circadian variability. Another important direction is to test whether the identified critical boundaries can predict seizure susceptibility under different rhythms of stimulation or pharmacological modulation. This would be particularly relevant for chronotherapy, where treatment timing could be optimized to keep the network farther from transition thresholds. More broadly, integrating patient-specific data with multiscale computational models may make it possible to identify individualized vulnerability windows and to develop more precise strategies for seizure forecasting and intervention.

Overall, our results support a dynamical view of epilepsy in which seizure susceptibility emerges from the interaction between pathological recruitment, intrinsic excitability, and circadian-like modulation of synaptic timing. Rather than acting as a direct trigger, physiological rhythm appears to shift the network across a landscape of critical boundaries that determine whether activity remains interictal or transitions into a persistent ictal state.

## 4 Methods

### 4.1 Network architecture and global measures

To investigate the emergence of pathological network dynamics, we constructed and analysed spiking neural networks composed of an inhibitory population of Fast Spiking (FS) neurons and excitatory populations of Regular Spiking (RS) and Impaired (IMP) neurons (Figure 1A). The networks consisted of 10 000 neurons, with 20% FS neurons and the remaining 80% excitatory neurons divided between RS and IMP populations in varying proportions to model different levels of pathological recruitment. Neurons were randomly connected with a connection probability of 5% and all populations received external stimulation driving network activity. The simulation protocol (detailed in Figure 1B) follows a three-state timeline: a 5-second transient phase, a 40-second stimulation period, and a 20-second recovery phase. To quantify the resulting dynamics, we use the mean firing rate and the Spike Tiling Coefficient (STTC) as a measure of synchrony (Figure 1C). Based on these measures, network activity was also classified into ictal and interictal regimes (see below for details).

### 4.2 Neuron models and Intrinsic Excitability

Individual FS and RS neurons were modeled using the Adaptive Exponential Integrate-and-Fire (AdEx) formalism [25, 26] while IMP neurons were described by an extended AdEx model including an additional slow variable (*z*) [15]. This variable phenomenologically captures ionic dysregulation and acts as a slow modulatory process controlling neuronal excitability.

For FS and RS neurons, the membrane potential *V* (*t*) and the adaptation current *w*(*t*) evolve according to:

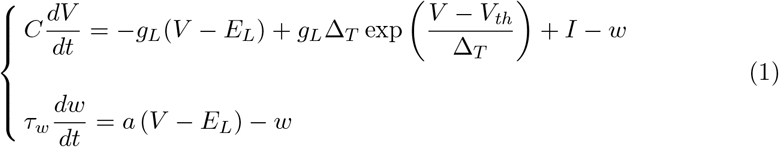

where *C* is the membrane capacitance, *g*_*l*_ the leak conductance, *E*_*L*_ the leak reversal potential, and Δ*T* controls the sharpness of spike initiation. The exponential term reproduces the rapid activation of sodium currents responsible for spike generation. The variable *w* represents spike-frequency adaptation, with *a* and *τ*_*w*_ controlling subthreshold coupling and adaptation timescale, respectively. *I* denotes the total synaptic and external input current. When the membrane potential crosses the spike threshold, the neuron emits a spike and the state variables are reset according to:

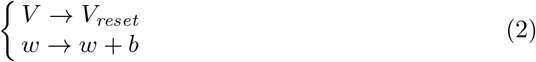

where *b* is the spike-triggered adaptation. As anticipated, IMP neurons were modelled using an extended AdEx formulation (as in [15]):

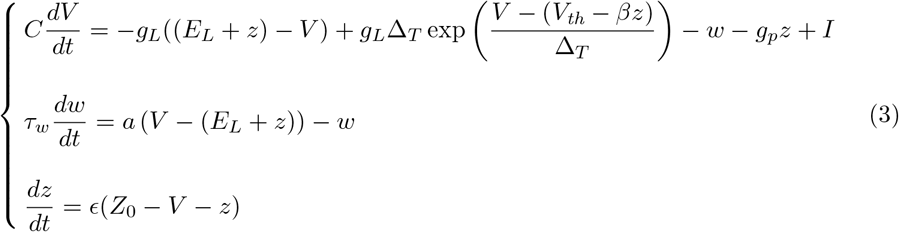

The slow variable *z* modulates both the effective resting potential and spike threshold, hence dynamically regulating neuronal excitability. This formulation is derived from a biophysical Hodgkin-Huxley-type description of excitability [27], in which variations in ionic gradients modify the membrane reversal potential through the Nernst relation. In the present model, *Z*_0_ controls the dynamics of *z* and therefore sets the intrinsic excitability level of the network. Physiologically, changes in *Z*_0_ can be interpreted as reflecting alterations in extracellular ionic concentractions of potassium (analogous the the [K]bath used in [27]). Increasing *Z*_0_ effectively depolarizes the neuron, bringing it closer to firing threshold and promoting seizure-like activity. Accordingly, *Z*_0_ was used as a control parameter to explore how intrinsic excitability shapes the dynamical regimes of the IMP population.

Single-neuron responses to current injection illustrate the distinct dynamics generated by the three neuron models. FS neurons displayed sustained high-frequency firing with no adaptation (Figure 2A and B, red), and RS neurons exhibited regular spiking with adaptation dynamics (Figure 2A and B, green). In contrast, IMP neurons showed progressive enhanced excitability, altered firing patterns, and slower cumulative dynamics as *Z*_0_ increases (Figure 2A and B, blue shades). These changes emerged from the slow accumulation of the variable, whose dynamics evolved over substantially longer timescales than the membrane potential and adaptation current, promoting prolonged depolarized states and sustained firing activity (Figure 2C). Increasing *Z*_0_ from *−*80mV to *−*10mV leads to a monotonic increase in firing frequency (Figure 2D). This transition was accompanied by larger steady-state values of, indicating that the slow modulatory process provides a persistent excitability drive that facilitates prolonged and high-frequency firing states.

### 4.3 Synaptic Dynamics

Synaptic interactions are mediated by conductance-based models, and hence the input current *I* is then defined as the sum of synaptic currents:

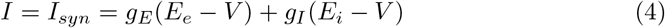

where *g*_*E*_ and *g*_*I*_ are the excitatory and inhibitory conductances, and *E*_*e*_ and *E*_*i*_ are the reversal potential of the excitatory and inhibitory synapses, respectively. The synaptic conductances are described by:

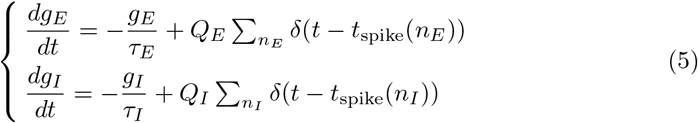

where *Q*_*E*_ and *Q*_*I*_ are the excitatory and inhibitory quantal conductances, *n*_*E*_ and *n*_*I*_ are the number of incoming spikes from excitatory and inhibitory synapses, *t*_*spike*_(*n*_*E*_) and *t*_*spike*_(*n*_*I*_) are the time at which the incoming excitatory and inhibitory spikes arrive, and finally *τ*_*E*_ and *τ*_*I*_ are the excitatory and inhibitory decay timescale.

To explore the impact of circadian modulation of synaptic parameters we define the synaptic time-scale ration as:

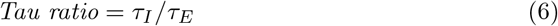

and systematically varied it to investigate its impact on the dynamics of the neural network. Specifically, the Tau ratio was explored over a range of values (between 0.8 and 1.2) covering physiologically plausible regimes, consistent with circadian modulations reported in the literature.

### 4.4 Population firing rates

To quantify the global activity of the different neuronal populations we computed the mean population firing rate over time. Spike trains, generated during the simulations, were binned in time and converted to firing rates by averaging across neurons within each population as follows:

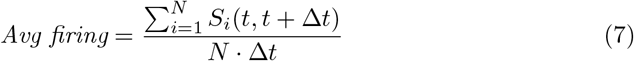

where *S*_*i*_ is the number of spikes from neuron *i* in the interval Δ*t*, and *N* is the total number of neurons in that specific population.

### 4.5 Network Synchrony (STTC)

Network synchrony was quantified using the Spike Time Tiling Coefficient (STTC), a pairwise measure of correlation between spike trains. The STTC between two neurons, *A* and *B* is defined as:

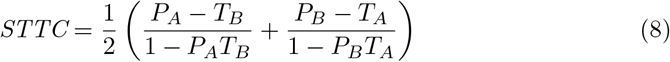

where *P*_*A*_ and *P*_*B*_ are the proportion of spikes in one train occurring within a temporal window *±*Δ*t* of spikes in the other train, while *T*_*A*_ and *T*_*B*_ are the fraction of the recording time that fall within *±*Δ*t* of spikes from each train. To capture the evolution of synchrony, we used a sliding window approach with a window size of 200ms. STTC values were computed between randomly sampled pairs of neurons within each population (corresponding to approximately 10% of the neurons in that population) and then averaged out to obtain a time-resolved synchrony estimate. In this way we obtained a continuous synchrony trace directly comparable with the evolution of the population firing rates and network state transitions. The STTC measure was chosen over other conventional measures because it effectively decouples temporal correlation from firing intensity, ensuring that the observed pathological synchronization reflects a reorganization of spike timing rather than a simple increase in population discharge frequency.

### 4.6 Ictal and Interictal regimes

An ictal state was defined as a sustained period (more than 500ms) during which the population firing rate exceeded a predefined threshold. The thresholds were defined according to the intrinsic firing properties of each population: 100Hz for FS and RS, and 125 Hz for IMP neurons. By contrast, interictal regimes were defined as periods in which the firing rate remained below 10Hz. And also in this case to avoid classifying brief fluctuations, only events lasting longer than 500ms were retained. Using these criteria, each simulation was analysed and the resulting classification was then combined with the STTC analysis to characterize how firing activity and synchrony co-evolved across different parameter regimes and levels of impaired neuronal recruitment.

## 5 Data availability

A subset of the synthetic data is available here: https://github.com/Computational-NeuroPSI/CMEI-Epilepsy.

In order to get access to the full dataset, please contact the corresponding author.

## 6 Code availability

The implemented Python and Brian 2 [28] codes are available here: https://github.com/Computational-NeuroPSI/CMEI-Epilepsy.

Simulations were performed on a Linux-based desktop computer equipped with an AMD Ryzen 9 3900X 12-core (24-thread) processor and 64 GB of RAM, and on high-performance computing (HPC) resources.

## 7 Acknowledgments

We thank Dr. Pascal Helson for the enlightening discussions. The computations were enabled by resources provided by the National Academic Infrastructure for Supercomputing in Sweden (NAISS) at PDC KTH, partially funded by the Swedish Research Council through grant agreement no. 2022-06725.

IC: Digital Futures. AD: Research supported by CNRS, Agence Nationale de la Recherche (FLAG-ERA BrainAct project, CR-CNS ImpactCom project), and the European Union (Human Brain Project H2020-945539 and Virtual Brain Twin project 101137289). DD and VJ: The preparation of this article was funded through the EU’s Horizon Europe Programme SGA 101147319 (EBRAINS 2.0), SGA 101137289 (Virtual Brain Twin), and No. 101057429 (project environMENTAL), and government grant managed by the Agence Nationale de la Recherche reference ANR-22-PESN-0012 (France 2030 program).

## 8 Author contributions

ND, IC, and DD conceived the study. ND and IC developed the model and methodology. IC performed the simulations, carried out the investigation, analysis, and prepared the visualizations. ND, IC, and DD wrote the original manuscript draft. AD and VJ contributed to review and editing. MHB supervised ND’s contribution, and DD supervised the project.

