## Supplementary Material for "Circadian Modulation of Excitation—Inhibition Balance Drives Ictal Transitions in a Mechanistic Model of Epileptic Networks"

### 1 Supplementary Material

The supplementary figures extend the main results by exploring additional parameter regimes and providing further evidence for the robustness of the reported transitions.

Supplementary Figure 1 illustrates how circadian fluctuations in the E/I balance modulate seizure susceptibility over time, highlighting how minima in the E/I ratio correspond to peaks in network vulnerability.

Supplementary Figures 2 and 3 extend the synchrony landscape analysis of the main text to additional IMP fractions and the FS population, characterizing how the asynchronous-to-synchronous transition and its persistence after stimulation offset depend on network composition and synaptic timescales.

#### 1.1 Supplementary Figures

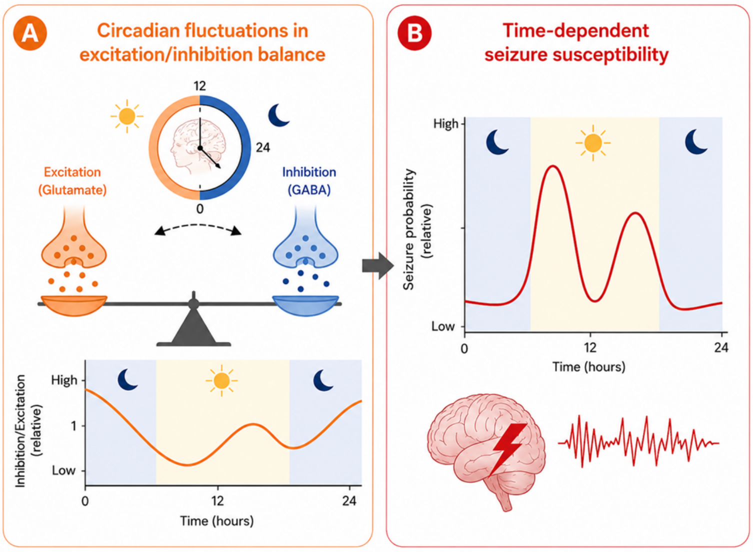


**Fig. 1 Circadian modulation of the excitation–inhibition balance and seizure susceptibility.** A) Schematic representation of circadian fluctuations in the excitation–inhibition (E/I) balance within cortical networks. Over the 24-hour cycle, coordinated variations in glutamatergic and GABAergic transmission dynamically modulate neuronal excitability. The lower panel illustrates the phenomenological evolution of the inhibition-to-excitation ratio, resulting from the interaction between a slow circadian component and secondary fluctuations associated with synaptic homeostatic regulation. Two relative minima emerge during the cycle, corresponding to transient states of increased network vulnerability. B) Conceptual framework linking the circadian dynamics of the E/I balance to seizure susceptibility. The red curve represents an idealized temporal profile of seizure occurrence probability, inspired by experimentally reported circadian distributions (adapted from [1]. The maxima of susceptibility coincides with the minima of the inhibigeneratedtion-to-excitation ratio observed in Panel (A), suggesting a mechanistic link between reduced inhibitory control, increased collective excitability, and the emergence of ictal transitions. This representation introduces the central hypothesis developed in the Results section: circadian rhythms may structure the dynamics of transitions.
The figure, originally generated in powerpoint, was subsequently refined using an artificial intelligence tool to improve its visual presentation.

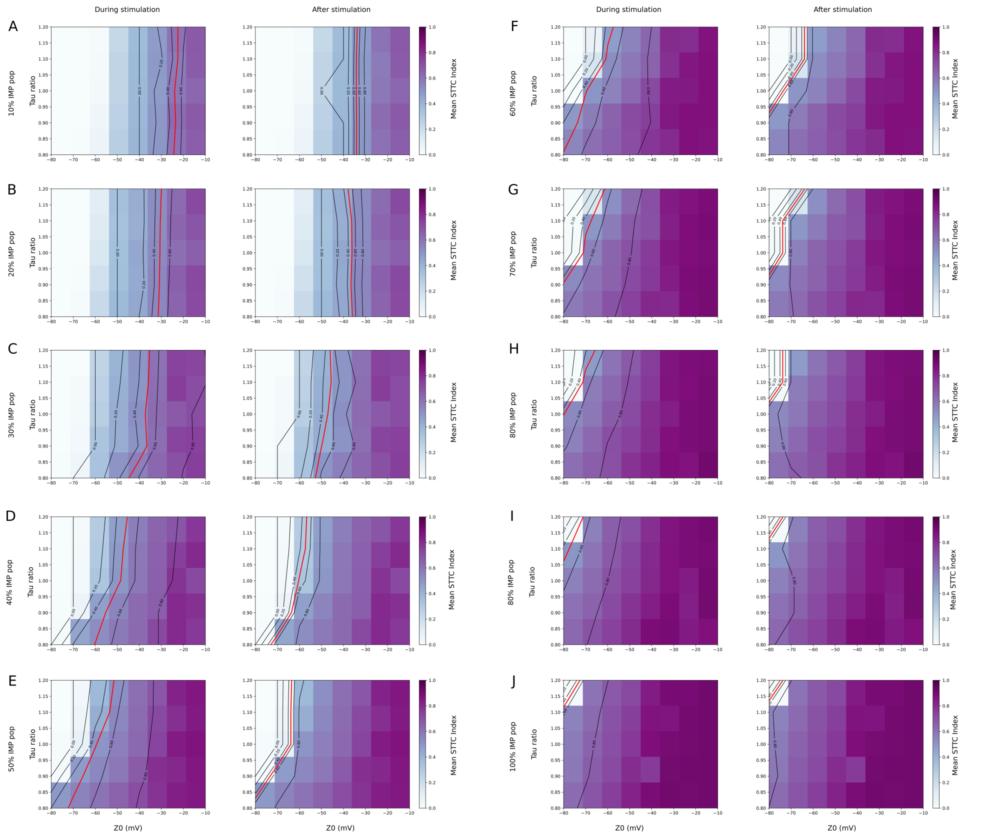


**Fig. 2 Synchrony landscape across parameter space and impaired population fraction in IMP population.** This figure is the expanded version of Figure 6. Here all the percentages of the IMP population are reported. Heatmaps show the mean Spike Time Tiling Coefficient (STTC), computed over IMP populations, as a function of the parameter $Z_0$ and the synaptic time-scale ratio (Tau ratio = $\tau_I$ / $\tau_E$), respectively on the x and y axis. Color indicates level of synchrony (mean STTC), black contour lines denote iso-levels of synchrony, with the red line highlighting the transition boundary (threshold=0.5) separating low- and high-synchrony regimes. Across all conditions, increasing Z0 leads to higher synchrony, but the structure of the transition depends strongly on the fraction of impaired neurons. At low IMP fractions (A-D), synchrony remains confined to a narrow region of parameter space with Tau ratio having just a minimal influence. At higher fractions (E-J) the high-synchrony regime expands, becoming more structured and is strongly modulated by the Tau ratio. Lower values (faster inhibitory dynamics compared to excitation) promote the emergence of synchrony at mode hyperpolarized $Z_0$, effectively shifting the transition boundary. Comparing stimulation and post-stimulation periods reveals how network dynamics evolve once the transient excitatory input stops, highlighting the persistence of synchronous activity in the absence of sustained drive.

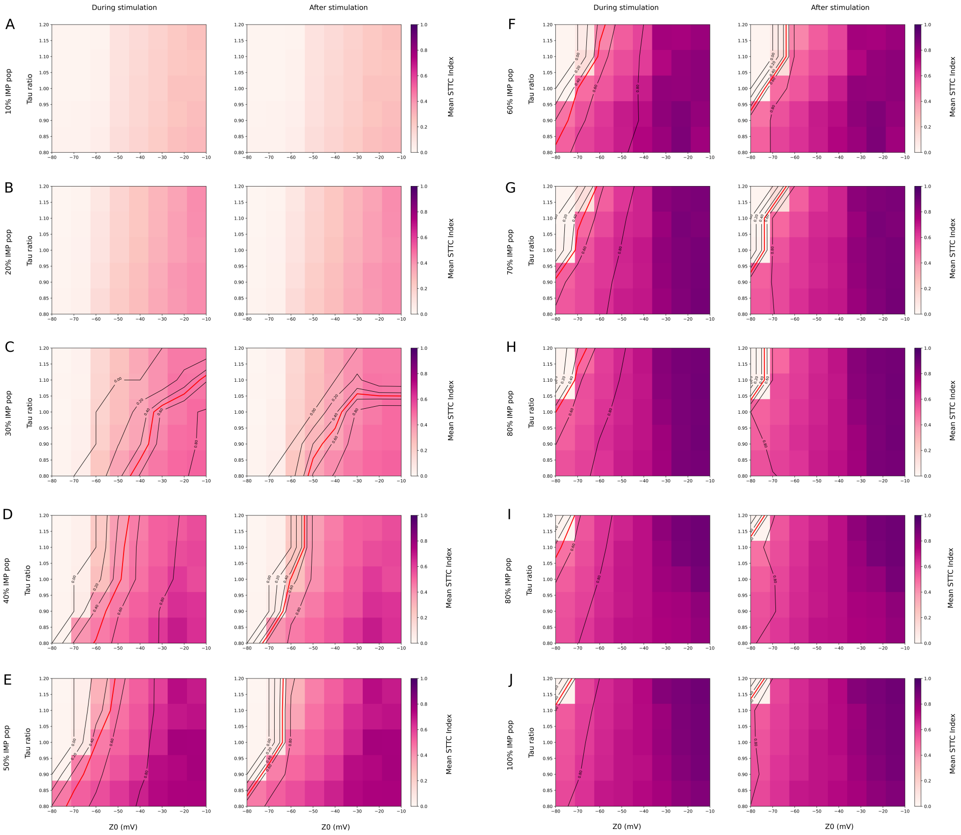


**Fig. 3 Synchrony landscape across parameter space and impaired population fraction in FS population.** Heatmaps show the mean Spike Time Tiling Coefficient (STTC), computed over FS populations, as a function of the parameter $Z_0$ and the synaptic time-scale ratio (Tau ratio $= \tau_I \ / \ \tau_E$), respectively on the x and y axis. Color indicates level of synchrony (mean STTC), black contour lines denote iso-levels of synchrony, with the red line highlighting the transition boundary (threshold=0.5) separating low- and high-synchrony regimes. Across all conditions, increasing $Z_0$ leads to higher synchrony, but the structure of the transition depends strongly on the fraction of impaired neurons. At low IMP fractions (A, B) the population does not reach the ictal state. As the IMP fraction increases (C-E) the FS population reaches the ictal regime and the Tau ratio has a big influence on the iso-levels. At higher fractions (F-J) lower values of Tau ratio (faster inhibitory dynamics compared to excitation) promote the emergence of synchrony at mode hyperpolarized $Z_0$, effectively shifting the transition boundary. Comparing stimulation and post-stimulation periods reveals how network dynamics evolve once the transient excitatory input stops, highlighting the persistence of synchronous activity in the absence of sustained drive. Interestingly, the iso-levels get closer to each other compared to the "during stimulation" counterparts
